# The 10,000 Immunomes Project: A resource for human immunology

**DOI:** 10.1101/180489

**Authors:** Kelly A. Zalocusky, Matthew J. Kan, Zicheng Hu, Patrick Dunn, Elizabeth Thomson, Jeffrey Wiser, Sanchita Bhattacharya, Atul J. Butte

## Abstract

New immunological assays now enable rich measurements of human immune function, but difficulty attaining enough measurements across sufficiently large and diverse cohorts has hindered describing normal human immune physiology on a large scale. Here we present the 10,000 Immunomes Project (10KIP), a diverse human immunology reference derived from over 44,000 individuals across 242 studies from ImmPort, a publicly available resource of raw immunology study data and protocols. We carefully curated datasets, aggregating subjects from healthy/control arms and harmonizing data across studies. We demonstrate 10KIP’s utility by describing variations in serum cytokines and leukocytes by age, race, and sex; defining a baseline cell-cytokine network; and using 10KIP as a common control to describe immunologic changes in pregnancy. Subject-level data is available for interactive visualization and download at http://10kImmunomes.org/. We believe 10KIP can serve as a common control cohort and will accelerate hypothesis generation by clinical and basic immunologists across diverse populations.

**One Sentence Summary:** An open online resource of human immunology data from more than 10,000 normal subjects including interactive data visualization and download enables a new look at immune system differences across age and sex, rapid hypothesis generation, and creation of custom control cohorts.

## Introduction

The rapid advancement of technologies in preclinical immunology (*1–5*) and the promise of precision therapeutics in immunology (*6–8*), have together propelled a rapid increase in the production of large-scale immunological data. Similar advancements in other fields, such as genomics, where high-throughput assays spurred a swell of data, have demonstrated the need and benefit of common reference datasets. Resources such as the 1000 Genomes Project (*9–11*), Health and Retirement Study (https://hrs.isr.umich.edu/), Wellcome Trust Case Control Consortium (*12*), and Exome Aggregation Consortium (*13*) have accelerated discovery of thousands of disease-linked variants and uniquely enable understanding of global variation in the human genome in health and disease. To date, however, human immunology has no such resource. A common reference would expand our understanding of the diversity of the human immune system, accelerate hypothesis testing, and serve as a common control population, precluding the need for immunologists to recruit such controls in every individual study.

The challenge in generating such a resource lies, in part, in the diversity of data types available to immunologists. A reference “immunome” might reasonably include flow cytometry, gene expression, human leukocyte antigen (HLA) type, cytokine measurements, clinical assessments, and more. Furthermore, standardized protocols for measurement and conventions for naming cell types and cytokines are only currently being developed, and adherence is inconsistent (*14*). For the experimental or clinical immunologist, the cost of generating the necessary data from scratch—or the temporal and computational costs associated with standardizing and harmonizing data from publicly available cohorts across platforms, time points, and institutions—is prohibitive. Thus, although the benefit of a common reference population is clear, and large-scale data are publicly available, this need has not been met.

Other lessons from the field of genomics offer additional direction and promise. For example, resources like the 1000 Genomes Project (*9–11*) have clearly demonstrated the necessity of exploring and accounting for human diversity; the publication of the original data release has been cited more than 5000 times. Additionally, although high-throughput assays invariably suffer from inter-experiment technical variation, the field has generated and validated statistical methods for overcoming those artifacts while preserving the underlying effects of interest (*15–19*). These breakthroughs, ripe for translation to immunological data, have unlocked the potential for deeper insight beyond the initial intent of each of the thousands of studies that have made their raw data publicly available to researchers.

Given the recent growth in open immunology data, we sought to synthetically construct a reference “immunome” by integrating individual level data from publicly available immunology studies, an effort which we term the 10,000 Immunomes Project (10KIP). We began by manually curating the entire public contents of ImmPort (Data Release 21; www.immport.org), the archival basic and clinical data repository and analysis platform for the National Institute for Allergy and Infectious Disease (NIAID) (*20,21*). ImmPort contains studies on a diversity of topics related to immunity, including allergy, transplant, vaccinology, and autoimmune disease, and the data represented are diverse, ranging from flow cytometry and Enzyme-Linked Immunosorbent Assay (ELISA) to clinical lab tests and HLA type. While most of these studies were not designed to examine the diversity of the healthy normal immune system, they nonetheless contain healthy control arms that we utilized for this purpose. ImmPort is uniquely suited to the task of generating a large diverse reference population of immune measurements, as the raw data deposition in ImmPort is highly structured. Every subject, sample, experiment, study, and experimental time point is assigned a unique accession, making every entity and attribute traceable throughout the database. Data contributors submit protocols detailing their process and the specific platforms and reagents used. Finally, every subject is associated with an age, sex, and race. This degree of annotation is rare amongst immune data repositories and is strictly necessary for compiling a diverse common reference population that enables custom cohort creation and systems-level analysis.

Our goal was to include in our reference only human subjects from the healthy control arms of studies and only samples from individuals that have undergone no experimental manipulation. Our filtering and data harmonization process resulted in an inaugural dataset consisting of 10 data types in standardized tables (mass cytometry [CyTOF], flow cytometry, multiplex ELISA, gene expression array, clinical lab tests and others) on 42,117 samples taken from 10,344 subjects. We unify the data from all the normal healthy immunomes into a fully open and interactive online resource (www.10kimmunomes.org). We expect that the ability to dynamically visualize and compare our reference population with samples from immune perturbation and disease will accelerate discovery in immunology. We further show that this resource can increase the potential for rapid hypothesis generation and testing, can serve as a common control population to increase the robustness of human immunology studies, and can also provide a basis for studying immunity across age, sex, and racially-diverse populations. As an arm of the ImmPort environment, the 10,000 Immunomes Project (10KIP) is highly scalable, and will only grow in value, richness, and scale with the participation of the immunology community in the open-data movement.

## Results

### Development of the 10,000 Immunomes Project

To develop the 10KIP, we began with ImmPort Data Release 21 (downloaded May 3, 2017), which contains 242 studies released to the public, with 44,775 subjects and 293,971 samples (fig. 1). We began by manually curating each of these 242 studies, reading inclusion and exclusion criteria, and selecting by hand which study arms and planned visits constitute data collected on samples from normal healthy human subjects prior to any experimental immune perturbation. This manual curation process resulted in an inaugural population of 10,344 subjects, spanning 83 studies and collectively contributing data from 42,117 biological samples. An exhaustive list of all studies, arms, and planned visits that qualified for inclusion is available as Table S1.

**Fig. 1.**
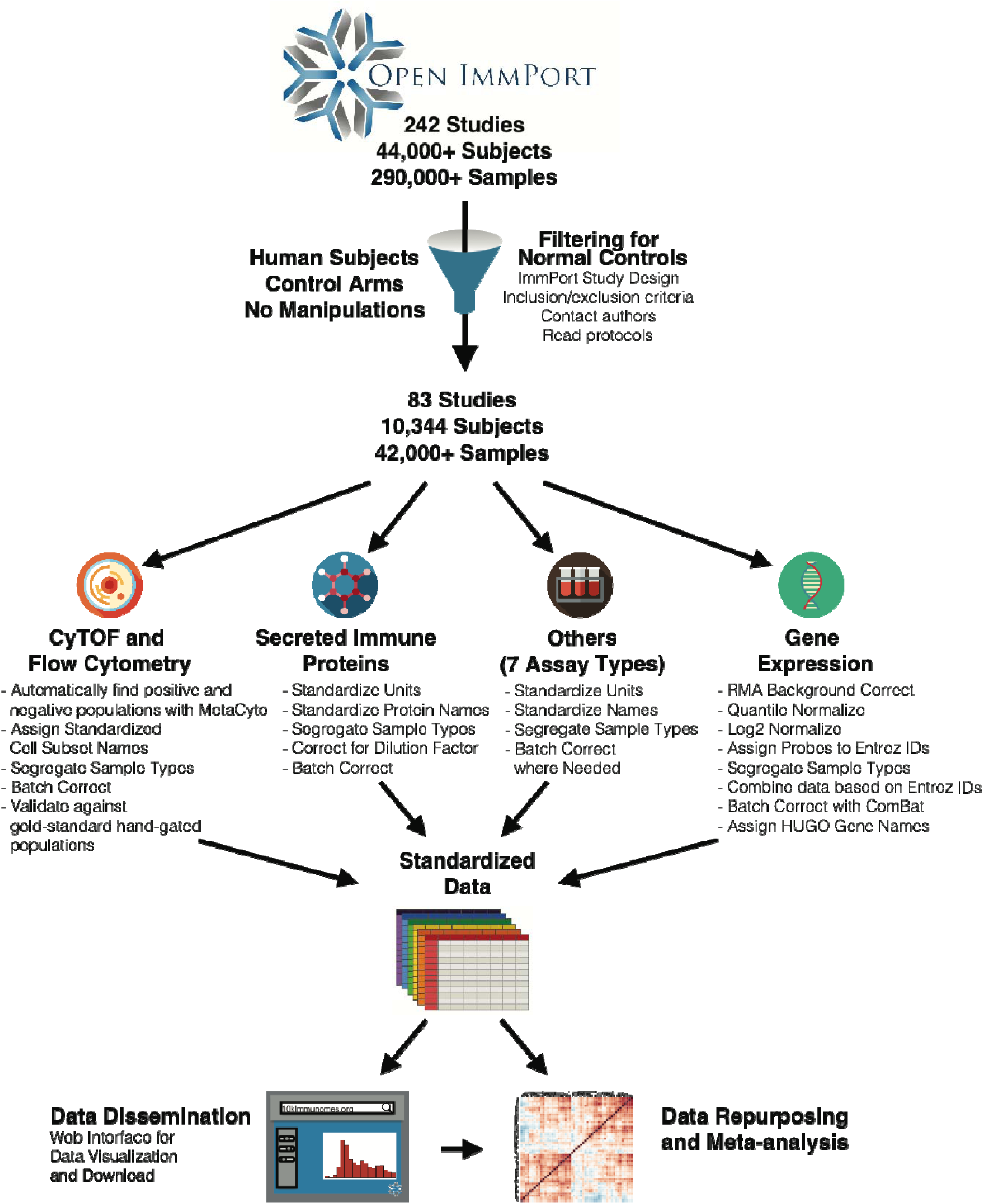
Resource Development and Selected Applications. Data from 242 studies and 44,775 subjects (including flow cytometry and CyTOF, mRNA expression, secreted protein levels—including cytokines, chemokines, and growth factors— clinical lab tests, HAI titers, HLA type and others) were collected from the NIAID Immunology Data and Analysis Portal, ImmPort (www.immport.org). We hand curated the entire contents of ImmPort to filter for normal healthy control human subjects. Each of the 10 data types was systematically processed and harmonized. These data constitute the largest compendium to date of cellular and molecular immune measurements on healthy normal human subjects. Both the normalized data and their raw counterparts are openly available for visualization and download at http://10kImmunomes.org/.

This dataset consists of 10 distinct data types (flow cytometry, high-throughput serum protein measurements, gene expression, clinical lab tests and others). For each data type, we developed a standardized pipeline for data cleaning and harmonization (see Methods). Across all studies, we standardized analyte names and units of measurement, segregated data by sample type (e.g. peripheral blood mononuclear cells (PBMC) versus whole blood versus serum), and corrected for differences in sample dilutions. This process resulted in standardized data tables, which form the backbone of the reference. The normalized data and their raw counterparts are available for visualization and download at http://10kImmunomes.org/.

### Contents of the inaugural release of the 10,000 Immunomes Project

The initial release of the 10KIP contains 10,344 subjects. They are approximately evenly split between male and female, represent a diverse racial makeup, and include more than 1,000 pediatric subjects (<18 years of age) and over 1,300 subjects above 65 years of age (fig. S1). As enumerated in Table 1, the resource contains secreted protein data from over 4,800 subjects, clinical lab test data from over 2,600 subjects, flow cytometry or mass cytometry data from over 1,400 subjects, HAI titers from over 1,300 subjects, and HLA types from over 1,000 subjects, in addition to several other data types. Because many subjects contribute more than one type of measurement, the total counts of subjects across all measurement types substantially exceeds the number of distinct subjects.

**Table 1.** Data Available in the Initial Release. Counts of distinct subjects for whom raw data of each type is represented in the initial release of the 10KIP. Because many subjects contributed multiple measurement types, the totals across all measurement types substantially exceed the number of distinct subjects.

| **Data available in the**<br>**10,000 Immunomes Project** |  |
| --- | --- |
| **Total Samples** | **42117** |
| **Total Distinct Subjects** | **10344** |
| **MEASUREMENT** | **SUBJECTS** |
| <u>**Secreted Proteins**</u> | 4835 |
| *ELISA* | 4035 |
| *Multiplex ELISA* | 1286 |
| <u>**Virus Titer**</u> | 3609 |
| *Virus Neutralization Titer* | 2265 |
| *HAI Titer* | 1344 |
| <u>**Clinical Lab Tests**</u> | 2639 |
| *Complete Blood Count* | 1684 |
| *Comprehensive Metabolic Panel* | 664 |
| *Fasting Lipid Profile* | 664 |
| <u>**Questionnaire**</u> | 1422 |
| <u>**Cytometry**</u> | 1415 |
| *Flow Cytometry (PBMC)* | 907 |
| *CyTOF (PBMC)* | 583 |
| *Flow Cytometry (Whole Blood)* | 164 |
| <u>**HLA Type**</u> | 1093 |
| <u>**Gene Expression Array**</u> | 476 |
| *Whole Blood* | 311 |
| *PBMC* | 165 |

### Multiplex ELISA measurements across the population

The regulation of immune system components through cytokines, chemokines, adhesion molecules, and growth factors is central to maintenance of a healthy immune homeostasis and response to acute infection (*22–25*). Recent advances in the measurement of such secreted proteins with multiplex ELISA (also known as multiplex bead-based analysis or by the trade name “Luminex”) allow for high-throughput profiling of the immune molecular milieu (*26,27*). Similar to high-throughput measurements of RNA expression, however, this type of measurement must be interpreted with caution, due to inter-experimental technical variation, as well as differences in reagents and platforms used (*28*). In fact, there is contention within the field of computational immunology regarding the validity of directly comparing high-throughput serum cytokine measurements across studies (*29,30*).

Here we suggest, however, that previously described models for statistical compensation for batch effects in genomics are sufficient for analysis of multiplex ELISA data. We find that, without batch correction, technical variation contributes significantly to the clustering of multiplex ELISA data as visualized by t-SNE (fig. 2A). The empirical Bayes algorithm ComBat (*15*), originally designed for analysis of microarray data, compensates for both mean and variance differences across studies while preserving potential effects of interest, such as differences by age, sex or race (fig. 2B, fig. S2). We have additionally confirmed the efficacy of this strategy through 1000-fold simulations of multiplex ELISA data with mean, variance, and single-analyte batch effects (fig. S2). This strategy preserves known effects, such as a significantly higher serum leptin concentration in women as compared to men (fig. 2C, (*31*)).

**Fig. 2.**
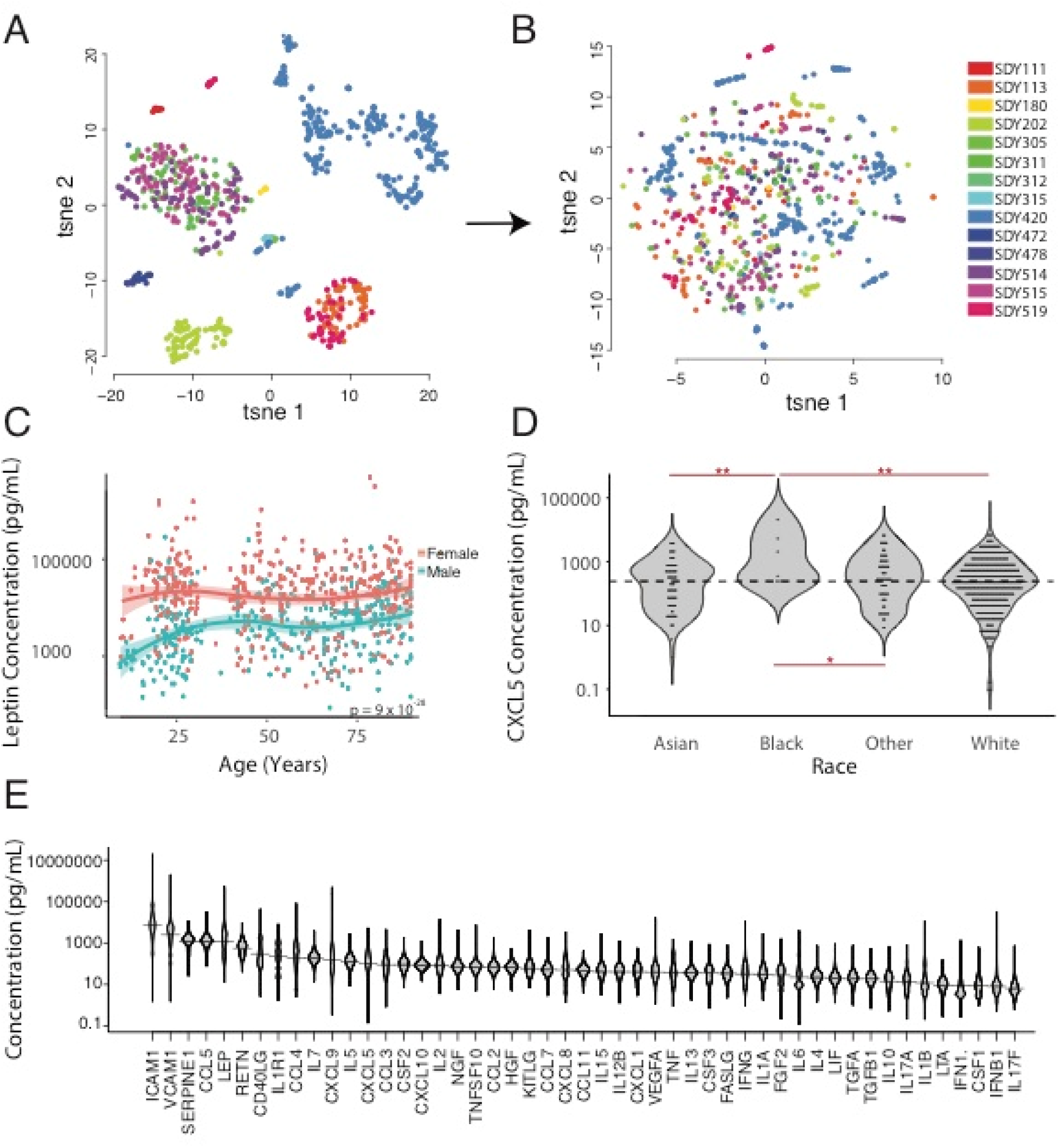
High-throughput secreted protein data: characterizing the range of unperturbed secreted protein levels in a diverse population. **(A)** tSNE visualization of high-throughput secreted protein data, colored by study accession, reveals that much of the variance across the data is explained by batch. **(B)** After batch correction with an empirical Bayes algorithm, which accounts for both mean and variance difference across studies while maintaining effects of covariates such as age, sex, and race, the data no longer cluster by batch. **(C)** Secreted protein data as measured by multiplex ELISA across 17 studies captures known effects, such as elevated levels of serum leptin in female relative to male subjects (ANCOVA, n = 906, p = 9 x 10^-28^). **(D)** Analysis of the reference population reveals novel demographic associations, including elevated CXCL5 in African American subjects as compared to other races. (ANCOVA, n = 917, p-values: * p < 0.05, ** p < 0.01). **(E)** We characterize the distribution of secreted protein levels from serum across the reference population (n = 1286).

Additionally, the ability to combine data across studies from disparate geographic locations and distinct ethnic populations enables us to find novel associations. Our analyses suggest, for example, a significantly higher level of C-X-C motif chemokine 5 (CXCL5) among African Americans as compared to other races (fig. 2D). This chemokine, which is produced by adipose tissue resident macrophages, is of potential clinical interest due to its association with insulin resistance (*32*). In total, we find that 27 out of the 50 most commonly measured cytokines, chemokines, and metabolic factors measured by multiplex ELISA differ significantly by age, sex, or race (fig. 2, S3). Finally, this analysis from a population of 1286 individuals across 17 studies allows us to describe the distribution of serum cytokine measurements in a diverse human population. Some, such as Interleukin-5 (IL5) and IL7, lie within a relatively small range, whereas others, such as chemokines C-C motif chemokine 4 (CCL4) and (CXCL9) display a many-fold range, even within this population of putative healthy normal human subjects. Together, these findings affirm the benefit of maintaining and growing a diverse common control population for the future of clinical and precision immunology.

### Individual level cell subset measurements across the population

Similarly, even within this reference population, we find a high degree of variability in the proportion of immune cell subsets from PBMC as measured by mass cytometry. This variability in cell subsets within a normal healthy population corroborates previously-reported descriptions of cell subset percentages (*33*). CD4^+^, CD8^+^, and gamma-delta T cell subsets, in particular, span a wide range as a percentage of total leukocytes. Smaller subsets, such as memory B cells and plasmablasts, span a tighter range (fig. 3A). For high-throughput analysis of mass cytometry data, we have employed a previously-validated pipeline that begins with raw fcs files, enacts quality control, implements automated gating based on a standard set of markers, and reports a standardized set of cell subset percentages as a proportion of total leukocytes (see Methods; (*34*)). In a prior publication, we enumerated a number of associations between race and cell-subset percentages from analysis of publicly available data (*34*).

**Fig. 3.**
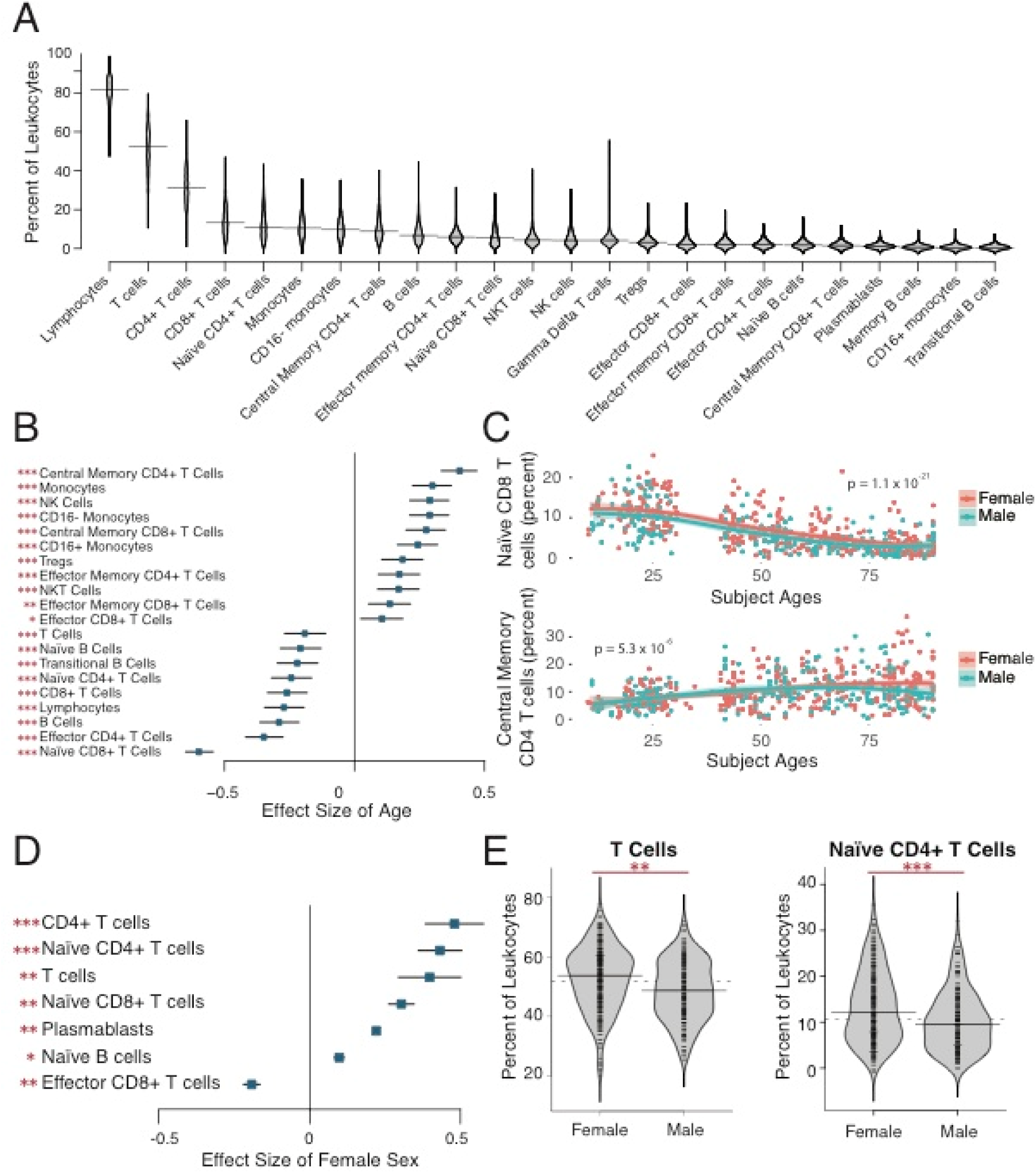
Mass cytometry: characterizing the range of cell subset percentages in a diverse population. **(A)** Distribution of cell subset percentages across the 10KIP. **(B)** Analysis of mass cytometry data reveals significant effects of age on cell subset percentages, while accounting for sex and race. Only cell subset associations with Benjamini-Hochberg corrected p-values < 0.05 are shown. (ANCOVA, n = 578, * p < 0.05, ** p < 0.01, *** p < 0.001). Effect sizes are displayed as Pearson’s r ± 95% confidence intervals. **(C)** Naïve CD8^+^ T cells decrease significantly with age (ANCOVA, n = 565, p = 1.1 x 10^-21^), while central memory CD4 T cells increase significantly with age (ANCOVA, n = 578, p = x 10^-6^), while accounting for sex and race. **(D)** Analysis of mass cytometry data reveals significant effects of sex on cell subset percentages, while accounting for age and race. Only cell subset associations with Benjamini-Hochberg corrected p-values < 0.05 are shown. (ANCOVA, n = 578, * p < 0.05, ** p < 0.01, *** p < 0.001). Effect sizes are displayed as Cohen’s d ± 95% confidence intervals. **(E)** T Cells (ANCOVA, n = 565, p = 7.4 x 10^-6^) and naïve CD4^+^ T cells (ANCOVA, n = 578, p = 3.3 x 10^-8^) are significantly elevated in women as compared to men, accounting for age and race.

Here, we describe the effects of age (fig. 3B, C), sex (fig. 3D, E), and race (fig. S4) in this larger healthy normal population. As an example, our analysis reveals a pronounced decline in naïve CD8^+^ T cells with age (fig. 3C) with a concomitant increase in memory CD4^+^ T cells (fig. 3C). These findings are anticipated given the accumulation of antigen exposures over the lifespan. Our analysis additionally suggests that women have significantly higher levels of naïve CD4^+^ T cells, naïve CD8^+^ T cells, naïve B cells, and plasmablasts than do male subjects, while having a significantly smaller proportion of effector CD8^+^ T cells (fig. 3D,E). We affirm the previously-published finding that Asian subjects have a lower proportion of CD4^+^ T cells as a proportion of leukocytes than do white subjects (*34*). We additionally find that NK cells are found at a significantly lower level in Asian subjects as compared to white subjects, and that regulatory T cells are measured at a significantly higher level in African American subjects as compared to all other races (fig. S4). These age, sex, and race-related differences in immune cell subsets may help explain population differences in infections and autoimmune disease or impact clinical decision-making as it pertains to treatment selection. Developing a diverse reference of immune measurements uniquely enables this type of discovery.

### Systems-level network analysis of cellular and molecular immunity

In addition to characterizing the diversity of the immune system in terms of cellular and molecular markers, the diversity of measurements available in the 10KIP also has the potential to facilitate systems-level network analysis. We selected 321 individuals from the dataset for which immune cell subsets in PBMC and protein measurements of serum cytokines, as measured respectively by mass cytometry and multiplex ELISA, were assessed in the same biological samples. We modeled the partial correlation between each cell type and each cytokine, statistically controlling for age, sex, and race (fig. 4), all of which our analyses suggest can have significant effects on cellular and molecular immune repertoire (fig. 2-3), and display only those correlations that remain significant at an FDR of 0.01 following Benjamini-Hochberg correction.

**Fig. 4.**
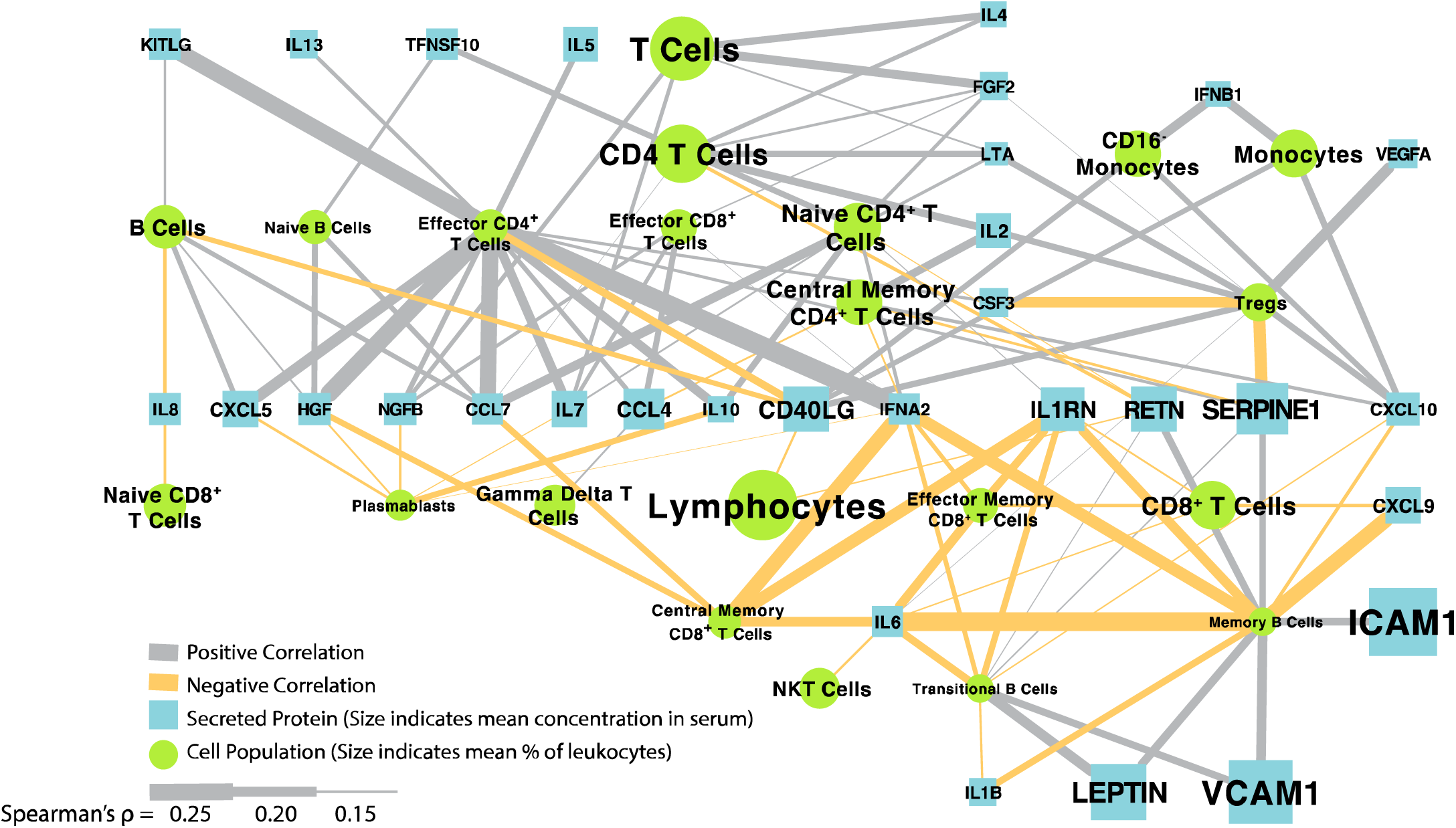
Immune cell and serum cytokine bipartite graph. Immune cell percentages and serum protein concentrations, as measured by CyTOF and multiplex ELISA, were processed as described in Methods, and the cell cytokine relationship was described as partial correlations accounting for age, sex, and race. Only relationships significant at a BH-corrected p < 0.01 are shown.

Our analysis recovers some known relationships; for example, we see that effector CD4^+^ T cells function as a major hub in the network, contributing positive associations with known Th2 cytokines IL5, IL10, and IL13 (*35*). We additionally see a negative association between regulatory T cells and the pro-inflammatory CSF3 (formerly granulocyte colony stimulating factor or GCSF)—consistent with the known immunomodulatory role of Treg cells (*36,37*). We detect an association between CXCL10 and monocyte subsets, concordant with evidence that this cytokine is expressed by and acts to recruit monocytes (*38*). Furthermore, acute phase reactants interferon alpha-2 and IL-6 are negative associated with central memory CD8+ T cells and memory B cells, which is concordant with the understanding of the kinetics of the transition from acute inflammation to memory formation (*39*). This exploratory analysis of the cell-cytokine network in the normal, healthy immune system also generates testable hypotheses about human immune function. For example, this analysis suggests a positive association between leptin and transitional and memory B cells, connections that are potentially of interest given B cell expression of the leptin receptor and the recent discovery that B cells may promote insulin resistance (*40,41*). Furthermore, this connection through memory B cells extends to the adipokine resistin and to the adhesion molecules ICAM-1 and VCAM-1, a cluster of molecules also known to be affected by adiposity (*42–44*). These analyses together demonstrate the utility of the 10KIP for generating systems-level hypotheses from large-scale publicly available immunology data collected for a variety of disparate purposes.

### Use as a common control population for precision immunology in pregnancy

Finally, to illustrate the potential of the 10KIP to serve as a common control group for clinical studies, we used an age and sex-matched subset of the 10KIP to compare with immune measurements in pregnancy, derived from ImmPort study SDY36. In this ImmPort study, researchers collected rich clinical data, as well as flow cytometry and serum cytokine measurements, from a population of 56 women during each trimester of pregnancy, six weeks postpartum, and six months postpartum. Cell count data from this study, as well as trends in cytokine secretion from cultured cells, have been published previously (*45*). Changes in serum cytokine levels over gestation and analyses of cell subset percentages (which are potentially differentially affected during pregnancy, (*46*)), however, remain undescribed. Additionally, the study design did not incorporate a pre-pregnancy control, leaving open the question of whether cell subsets and serum cytokines truly return to baseline by six months postpartum. Given work demonstrating persistence of fetal cells and DNA in maternal blood and brain many years postpartum (*47,48*), the comparison to a common control has the potential to enrich our understanding of the immune system in pregnancy and maternity.

We first applied principal components analysis (PCA) to the serum cytokine measurements, which revealed a major shift in cytokine regulation during the first trimester of pregnancy as compared to second and third trimester measurements, postpartum measurements, and measurements taken from age and sex-matched 10KIP controls (fig. 5A). This shift is primarily driven by increased concentrations of CCL2, CCL3, CCL4, CCL5, CCL11, CXCL10, and IL6. As an example of this modulation, we see that CCL5 concentration is significantly increased during the first and second trimester, decreased during the third trimester and up to 6 weeks postpartum, but returns to baseline by 6 months postpartum (fig. 5B). In contrast, IL15 measurements remain relatively constant over the entire course of gestation (fig. 5C).

**Fig. 5.**
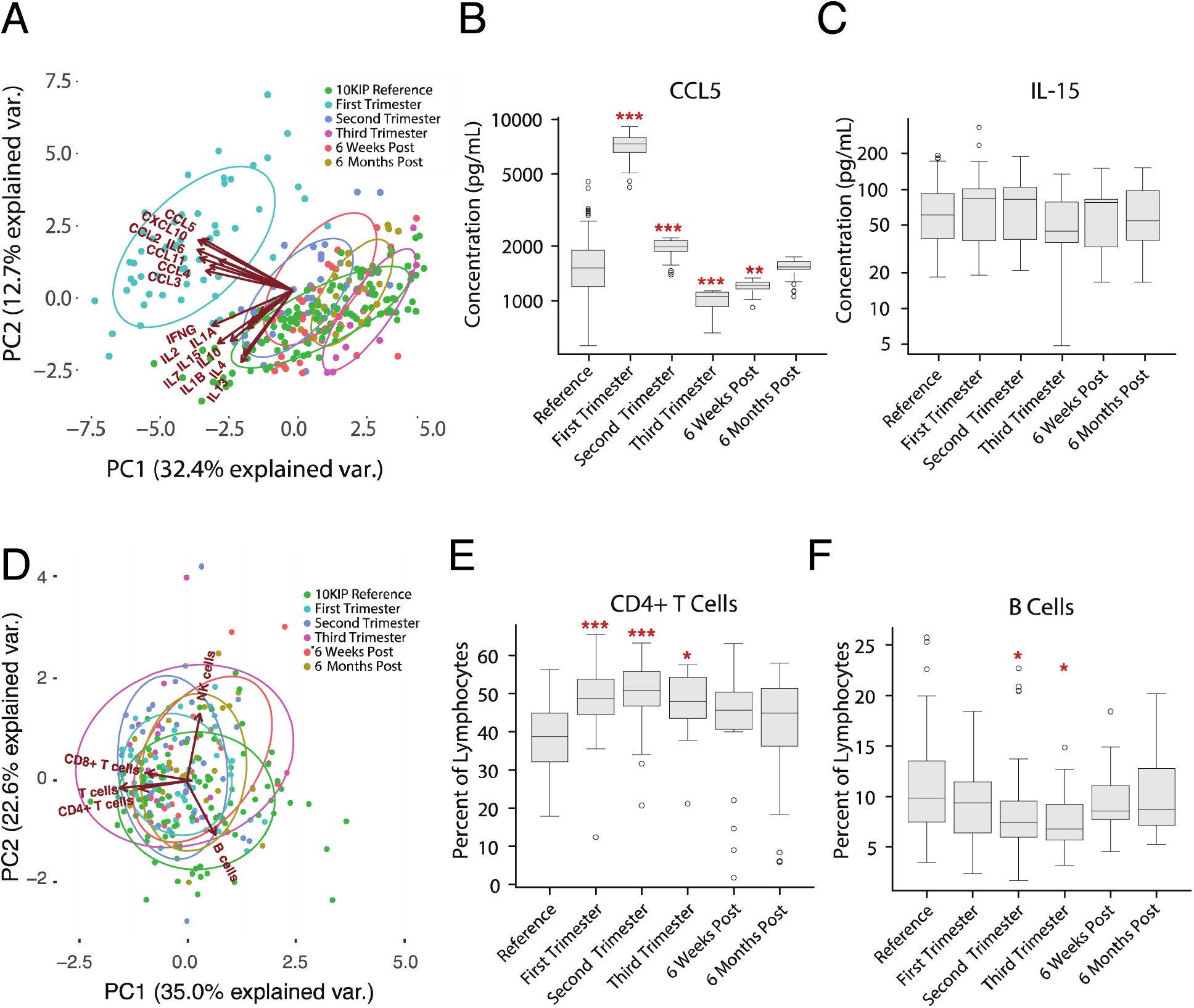
Comparing pregnancy data to the common control reveals cell-subset and immune protein modulation in pregnancy. **(A)** PCA plot depicting the variation in serum proteins, as measured by multiplex ELISA, over the course of pregnancy, taken from ImmPort Study SDY36, as compared to multiplex ELISA measurements from women between the ages of 18-40 from the reference population. The variance in measurements is dominated by a deviation in serum cytokine measurements during the first trimester (teal) relative to all other time points during pregnancy and relative to the 10KIP controls (green). These differences are driven primarily by changes in CCL2, CCL3, CCL4, CCL5, CCL11, IL6, and CXCL10. **(B)** As an example of cytokine modulation in pregnancy, serum CCL5 levels are significantly increased in the first and second trimester relative to the 10KIP controls, decrease during the third trimester and remain low for at least 6 weeks postpartum. CCL5 levels return to baseline levels by 6 months postpartum (ANOVA with Tukey HSD, n = 142 controls, n = 57 pregnancy, * p < 0.05, ** p < 0.01, *** p < 0.001). **(C)** In contrast, serum IL15 levels make no significant deviations from normal over the course of pregnancy (ANOVA with Tukey HSD, n = 142 controls, n = 57 pregnancy). **(D)** PCA plot depicting the variation in immune cell subsets, as measured by flow cytometry, over the course of pregnancy, taken from ImmPort Study SDY36, as compared to cytometry measurements from women between the ages of 18-40 from the 10KIP controls. As opposed to cytokine measurements (A), the preponderance of variation in cell subset measurements is not due to changes over the course of pregnancy. All time points during and following gestation substantially overlap with the controls (green). **(E)** The percentage of CD4^+^ T cells, as a fraction of lymphocytes, is significantly elevated over the duration of pregnancy, but returns to baseline in the postpartum period (ANOVA with Tukey HSD, n = 94 controls, n = 57 pregnancy, * p < 0.05, *** p < 0.001). **(F)** The percentage of B cells, as a fraction of lymphocytes, exhibits a small but significant dip in the second and third trimesters (ANOVA with Tukey HSD, n = 94 controls, n = 57 pregnancy, * p < 0.05).

In addition to analysis of serum cytokine concentrations, we also examined changes in cell subset percentages in pregnancy. PCA analysis of flow cytometry measurements indicated that changes in cell subsets over the course of gestation are not the primary source of variation as compared to postpartum or reference measurements (fig. 5D). This is not to say, however, that cell subsets remain static over the course of pregnancy. We see, for example, that CD4^+^ T cells, as a percent of lymphocytes, undergo a significant increase during all three trimesters of pregnancy as compared to the 10KIP reference population (fig. 5E). B cells, in contrast, exhibit a small but significant dip during the second and third trimesters (fig. 5F). This analysis demonstrates that the size and scope of the 10KIP are sufficient to generate age and sex-matched control cohorts for two types of high-throughput immune measurements as a baseline or comparator to immune perturbation or disease. In clinical studies, where challenges of compliance and recontactability limit researchers’ ability to generate well-powered control cohorts, common reference populations such as the 10KIP could accelerate discovery and increase confidence in findings of potential clinical import.

## Discussion

Although the availability of large common control cohorts, such as the 1000 Genomes Project (*9–11*) and the Wellcome Trust Case Control Consortium (*12*) has proven immensely useful for various biological research communities, no parallel resource exists for immunological measurements. Here we produced, through manual curation and study-by-study harmonization, the 10,000 Immunomes Project, a standardized reference dataset for the immunology community. To enable its use by experimental and clinical immunologists, we developed a framework for interactive data visualization, as well as custom cohort creation and data download, available at http://10kImmunomes.org/. Through statistical testing and validations in simulated data, we demonstrate the ability to compensate for technical artifacts that invariably arise from collecting data on different days, across different platforms, or at distant institutions, by repurposing algorithms developed in computational genetics.

In doing so, we recover known differences by age and sex across serum cytokine and cell-subset measurements, but also reveal differences, particularly by race, that would have been impossible to uncover without the combination of dozens of independent datasets generated to answer varied and unrelated questions in immunology. Through network analysis, we additionally demonstrate the utility of the resource for generating insights into cell-cytokine relationships in the human immune system. Finally, we demonstrate that the size and scope of the data are sufficient for custom cohort selection, enabling us to generate a reference cohort of women between the ages of 18 and 40 who have both cell subset and serum cytokine data available on the same blood samples for comparison with an external dataset derived from measurements taken during pregnancy. Generating a sufficiently powered sex and age-matched population with multiple immune measurements for comparison allowed us to explore the cell-subset and cytokine changes that occur as the immune system is modulated over the course of gestation. We expect that the 10KIP will prove useful as a common reference population for a diversity of future clinical and preclinical immunological studies.

Here, we demonstrate that the abundance of well-documented publicly available immune measurements in ImmPort is now sufficient to create an open data platform for human immunology. While we recognize the ideal would be to recruit and collect immune measurements from a large cohort, the resources required to collect immunologic measurements from a sufficiently sized heterogeneous population would be considerable, and a sizable volume of subject-level human immunology data are currently available to the research community. We also acknowledge further potential limitations in this work. For example, we are selecting subjects, standardizing labels and units, and otherwise curating the data with the best available information on these studies, but it is possible errors in the original data descriptions or labeling might persist. Also, we present the data after normalization and batch correction, but of course, we recognize that all of these source data sets were collected independently across institutions, technologies, and time. It is possible that our normalization efforts and assumptions might not hold true for every analysis in every study. We also note that, although the cohort is large and will continue to grow, some data types might not be measured densely enough to make reliable models that span all ages or races, and to date, racial information in ImmPort is acquired at a relatively coarse grain. As high-throughput immunological techniques become more widely available and as experimentalists continue to deposit these data in ImmPort, however, the resource will continue to grow, enabling well-powered analyses on more specific populations and over an increasing number of data types with time.

Finally, we want to recognize current reference datasets for immunology. The extant resources, while of clear import to the research community, serve different purposes than does the 10KIP. ImmGen (*49*), for example, represents an immense resource of immune gene expression in murine models, while the 10KIP instead focuses on multiple data types in human immunology. Likewise, ImmuneSpace (*50*) provides a suite of visualization and analysis tools, allowing users to interact and download data at the level of individual human immunology studies. The 10KIP, in contrast, has as its primary goals to filter the extant data for only healthy normal subjects, and to enable visualization and analysis across many studies. The 10KIP takes full advantage of the structure of ImmPort, in which subjects are assigned a unique accession number and are associated with their age, sex, and race. The resource allows researchers to subset the population or to look for associations with these general demographic phenotypes. Additionally, it leverages the richness of data available through ImmPort, which encompasses soluble protein and cytokine measurements, such as multiplex ELISA, cell-phenotyping measurements such as flow-cytometry and CyTOF, standard medical laboratory test panels, gene expression data, and others. We believe that integrating these datasets and presenting them as a fully open resource will pay dividends in terms of both basic research and the precision and robustness of ongoing translational efforts in immunology.

## Materials and Methods

### Subjects

Subjects were extracted from Data Release 21 of ImmPort database, which contains 242 open-access studies, together comprising 44,775 subjects and 293,971 samples. Each subject in ImmPort is assigned a unique identifier, allowing every measurement in the ImmPort database to be assigned to a unique subject. Each subject has, at minimum, race, age, and sex demographic information. The ImmPort data architecture requires that each study contain detailed descriptions of inclusion and exclusion criteria for subjects. Additionally, each arm (experimental and control arms) of each study is assigned a unique accession. Finally, each experimental measurement is time stamped with a unique planned visit accession. Manual review of the inclusion/exclusion criteria, arms, and planned visits allowed us to select control subjects, and to examine only those measurements taken before the onset of any experimental manipulation, such as vaccine, drug, or surgery that may have occurred. A complete list of qualifying studies, arms, and planned visits contained in the 10KIP is available in Table S1.

### Extract Immune Cell Frequencies from Cytometry Data

Meta-analysis of Cytometry data is conducted using the MetaCyto package (*34*). Briefly, flow cytometry data and CyTOF data of healthy human blood samples from ImmPort studies SDY89, SDY112, SDY113, SDY144, SDY167, SDY180, SDY202, SDY212, SDY296, SDY305, SDY311, SDY312, SDY314, SDY315, SDY364, SDY368, SDY387, SDY404, SDY420, SDY472, SDY475, SDY478, SDY514, SDY515, SDY519, SDY702, SDY720, and SDY736 were downloaded from ImmPort web portal. Flow cytometry data from ImmPort were compensated for fluorescence spillovers using the compensation matrix supplied in each fcs file. All data from ImmPort were arcsinh transformed. For flow cytometry data, the formula f(x) = arcsinh (x/150) was used. For CyTOF data, the formula f(x) = arcsinh (x/8) was used. Transformation and compensation were done using the *preprocessing.batch* function in MetaCyto (*34*). The cell definitions from the Human ImmunoPhenotyping Consortium (*14*) were used to identify 24 types of immune cells using the *searchClster.batch* function in MetaCyto. Specifically, each marker in each cytometry panels was bisected into positive and negative regions. Cells fulfilling the cell definitions are identified. For example, the CD14^+^ CD33^+^ CD16^-^ (CD16^-^ monocytes) cell subset corresponds to the cells that fall into the CD14^+^ region, CD33^+^ region and CD16^-^ region concurrently. The proportion of each cell subsets in the PBMC or whole blood were then calculated by dividing the number of cells in a subset by the total number of cells in the blood. Differences by age, sex, and race were detected with a linear model, with Tukey’s Honestly Significant Difference (Tukey’s HSD) post-hoc tests and Benjamini-Hochberg correction for false discovery rate.

### Multiplex ELISA analysis

Secreted protein data measured on the multiplex ELISA platform were collected from ImmPort studies SDY22, SDY23, SDY111, SDY113, SDY180, SDY202, SDY305, SDY311, SDY312, SDY315, SDY420, SDY472, SDY478, SDY514, SDY515, SDY519, and SDY720. Data were drawn from the ImmPort parsed data tables using RMySQL or loaded into R from user-submitted unparsed data tables. We corrected for differences in dilution factor and units of measure across experiments and standardized labels associated with each protein as HUGO gene symbols. This step represents the “formatted” multiplex ELISA data table. For our own analysis, as represented in Figure 2, we analyzed only those proteins that were measured in more than half of the subjects leaving the 50 most-commonly measured proteins. Compensation for batch effects was conducted using the *ComBat* function of the R package *sva*, with study accession representing batch and a model matrix that included age, sex, and race of each subject. Data were log2 transformed before normalization with ComBat to better fit the assumption that the data are normally distributed. We verified that a linear model associating age, sex, ethnicity, and study accession of each subject no longer revealed any significant associations between study accession and protein concentration following batch correction, and that known differences, such as the difference in leptin concentration by sex, were captured following our batch correction procedure. We additionally validated our approach using 1000-fold data simulations (see below). Differences by age, sex, and race were detected with a linear model, with Tukey’s Honestly Significant Difference post-hoc tests and Benjamini-Hochberg correction for false discovery rate.

### Network Analysis

The bipartite network depicted in Figure 4 represents an analysis over the 24 immune cell subset percentages calculated in the mass cytometry analysis described above in *Extract Immune Cell Frequencies from Cytometry Data* and the 50 soluble protein measurements, normalized and batch-corrected as described above in *Multiplex ELISA analysis*. Data were included from the 321 subjects where both multiplex ELISA and mass cytometry measurements were conducted on the same biological sample. Edges depict the Spearman’s ρ of a partial correlation between each cytokine concentration and each individual cell type, accounting for age, sex, and race. Only correlations that remained significant at a BH-corrected p < 0.01 are shown.

### Cell and cytokine modulation in pregnancy

We compared serum cytokine and cell subset percentages from 10KIP samples to measurements taken from women during and after pregnancy. We selected samples from the 10KIP from women aged 18-40 who contributed CyTOF data from PBMC and multiplex ELISA measurements. Samples from pregnancy were taken from ImmPort study SDY36. The serum cytokine and flow cytometry from SDY36 was batch corrected together with the ImmPort reference data, using the default parameters of the ComBat algorithm, and including age, sex, race, and time point in pregnancy in the model while using study accession as a surrogate for batch. Because SDY36 measured a smaller number of cytokines and cell subsets than are available as part of the 10KIP, we further selected a subset of the 10KIP to include just those parameters measured in SDY36. These data were used to conduct standard PCA analysis (R: prcomp, ggbiplot). Differences were calculated using ANOVA with a Tukey’s HSD post-hoc test.

### Gene expression array harmonization and normalization

Gene expression array data were obtained in three formats. For data collected on Affymetrix platforms, we utilized the *ReadAffy* utility in the *affy* Bioconductor package to read in raw .CEL files. The *rma* utility was used to conduct Robust Multichip Average (rma) background correction (as in (*51*)), quantile normalization, and log2 normalization of the data. For data collected on Illumina platforms and stored in the Gene Expression Omnibus (GEO) database, we utilized the *getGEO* utility in the *GEOquery* Bioconductor package to read the expression files and the *preprocessCore* package to conduction rma background correction, quantile normalization, and log2 normalization of the gene expression data. Finally, for data collected on Illumina platforms but not stored in GEO, we utilized the *read.ilmn* utility of the *limma* Bioconductor package to read in the data, and the *neqc* function to rma background correct, quantile normalize, and log2 normalize the gene expression data. In all instances, probe IDs were converted to Entrez Gene IDs. Where multiple probes mapped to the same Entrez Gene ID, the median value across probes was used to represent the expression value of the corresponding gene. The background-corrected and normalized datasets were combined based on common Entrez IDs, missing values were imputed with a k-nearest neighbors algorithm (R package: impute, function: impute.knn) using k = 10 and default values for rowmax, colmax, and maxp. To create the normalized and batch corrected dataset available through the www.10kImmunomes.org portal, we utilized a well-established empirical Bayes algorithm for batch correction (*15*), compensating for possible batch effects while maintaining potential effects of age, race, and sex across datasets and mapped Entrez IDs to HUGO gene IDs.

### Simulations to validate the batch correction algorithm

The empirical Bayes algorithm we have used to generate the normalized data available for download has previously been validated in its use for gene expression microarray analysis (*15*). To assess the efficacy of using an empirical Bayes algorithm to compensate for batch effects in multiplex ELISA data, we generated simulated multiplex ELISA data as skewed normal distributions from a set of parameters selected to mimic those skewed normal distributions that best fit the actual multiplex ELISA data used in our analysis. We generated this data for 50 analytes and 1500 subjects and purposefully introduced batch effects intended to mimic the types of batch effects we might encounter in real multiplex ELISA data. To account for use of a differently calibrated machine, for example, we simulated data in which one batch had a higher mean that the other batches. To account for the possibility that one lab’s data might be more variable than others, in one simulation we introduced random noise into one batch of the data. Finally, to account for the fact that the antibodies used may differ in efficacy across lots and experiments, we devised a simulation in which just one analyte in just one batch has a perturbed mean. In each of 1000 simulations of this data, we then generated a linear model to test whether the empirical Bayes algorithm ComBat (*15*) would successfully correct for these deviations from the true value of the simulated data. Additionally, we took the largest single batch of multiplex ELISA data (data from ImmPort study SDY 420) and intentionally introduced the same 3 types of batch effects we introduced into the simulated data. Following the same procedure, we demonstrate that ComBat successfully removes these introduced batch effects from real multiplex ELISA data.

## Acknowledgments

We thank M. Sirota, H. Maecker, Y. Rosenberg-Hasson, M. Spitzer, and J. Puck for their guidance and early advice in this work. We would also like to acknowledge all of the original data contributors, who by submitting their individual level data to ImmPort made this work possible. This work was supported by the National Institute of Allergy and Infectious Diseases (Bioinformatics Support Contract HHSN272201200028C). The content is solely the responsibility of the authors and does not necessarily represent the official views of the National Institutes of Health. KAZ led the design and implementation of the work, in collaboration with ZH for analysis of cytometry data, with PD, ET, and JW for building and curation of the ImmPort database, and with MJK, SB, and AJB for continual feedback regarding design and analysis throughout the project. The paper was written by KAZ with editorial input from all authors. The authors declare no competing interests. All data are available at http://10kImmunomes.org/.

## Supplementary Materials

**Fig. S1.**
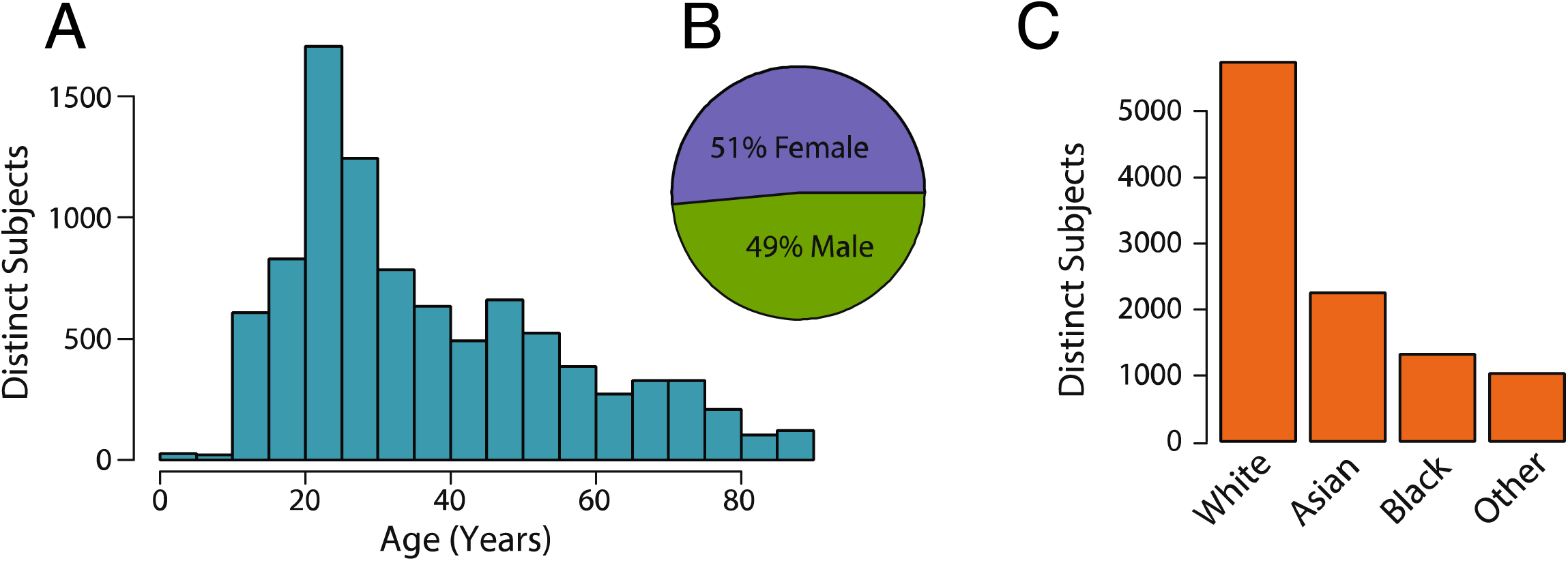
Demographics of the reference population. **(A)** Age distribution of the reference population. **(B)** Sex distribution of the reference population. **(C)** Racial distribution of the reference population.

**Fig. S2.**
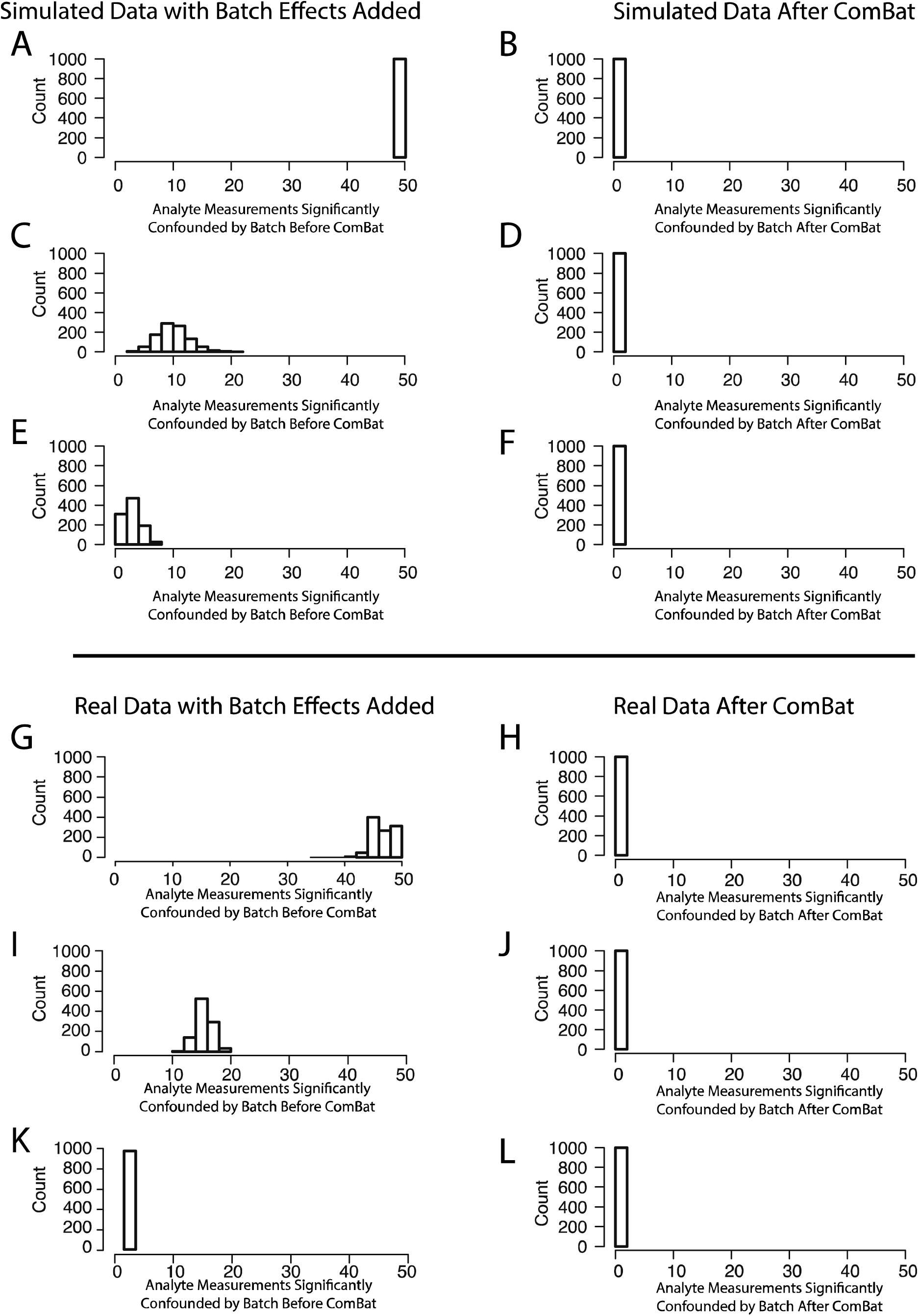
Batch correction of multiplex ELISA data. Because the linear model tests the effect of batch on each of the 50 analytes independently, the output of each run of the simulation could vary between 0 and 50 significant batch-analyte associations. Results are reported as mean number of analytes that are significantly affected by batch in a linear model ± the standard deviation in the number of analytes over the 1000 iterations. (**A-F**) 1000 different simulated multiplex ELISA datasets were generated as described in the Methods. In each of these 1000 simulations, three conditions were tested. 1) One entire batch has an increased mean value. 2) One entire batch has increased variance. 3) One analyte in one batch has a decreased mean value. For each condition, the data were altered to add this batch effect, and the data were then processed with ComBat to attempt to remove this batch effect. **A,C,E** represent the uncorrected data. **B,D,F** represent data after batch correction. **(A)** Over the 1000 runs of the simulation there were, in the mean perturbed data, 50 significant batch-analyte associations in every run. **(B)** This was reduced to 0 ± .08 following batch correction. **(C)** In the variance-perturbed data, there were 10 ± 2.7 significant batch analyte associations. **(D)** This was reduced to 0.6 ± 0.5 following batch correction. **(E)** In the single analyte perturbed data, there were 3 ± 1.5 significant batch-analyte associations (an increase of 1 over baseline) **(F)** This was reduced to 0 ± 0.04 following batch correction. These findings, together with our ability to recover known effects of demographic variables in multiplex ELISA data, provided confidence that the Empirical Bayes method of compensating for batch effects was reasonable and effective for the data at hand. (**G-L**) We also selected the largest single batch of multiplex ELISA data, data taken from ImmPort study SDY420, and tested the same three conditions as above in a 1000-fold simulation. **G,I,K** represent the uncorrected data. **H,J,L** represent data after batch correction. **(G)** Over the 1000 runs of the simulation there were, in the mean perturbed data, 47 ± 1.8 significant batch analyte associations. **(H)** This was reduced to 0 ± 0 following batch correction. **(I)** In the variance-perturbed data, there were 16 ± 1.6 significant batch analyte associations. **(J)** This was reduced to 0 ± 0 following batch correction. **(K)** In the single analyte perturbed data, there were 3 ± 0.04 significant batch-analyte associations (an increase of 1 over baseline) **(L)** This was reduced to 1 ± 0.13 following batch correction. These findings, together with our ability to recover known effects of demographic variables in multiplex ELISA data, provided confidence that the Empirical Bayes method of compensating for batch effects was reasonable and effective for the data at hand.

**Fig. S3.**
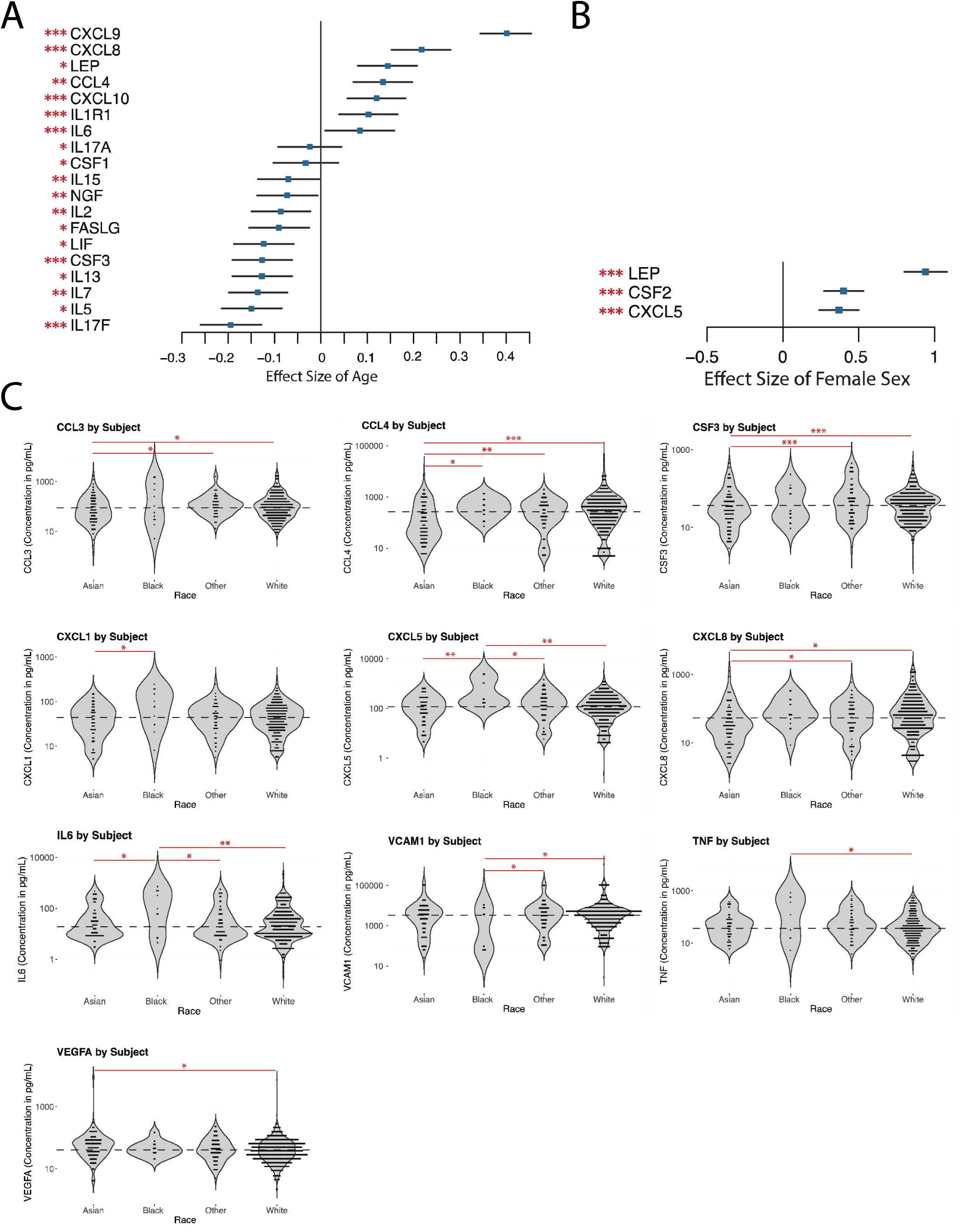
Multiplex ELISA measurements differ by age, race, and sex. **(A)** Multiplex ELISA data were processed as described in Methods. 20 out of the 50 most commonly measured serum proteins were significantly associated with age. Only proteins significantly associated with age at a threshold of BH-corrected p < 0.05 are shown. (ANCOVA, n = 1286, *** p < 0.001, ** p < 0.01, * p < 0.05). Effect sizes are displayed as Pearson’s r ± 95% confidence intervals. **(B)** Leptin, CXCL5, and CSF2 are all expressed at significantly lower levels in men as compared to women. Only proteins significantly associated with age at a threshold of BH-corrected p < 0.05 are shown. (ANCOVA with Tukey HSD, *** p < 0.001). Effect sizes are displayed as Cohen’s d ± 95% confidence intervals. **(C)** We found 10 of the 50 most commonly measured serum proteins to differ significantly by race (ANCOVA with Tukey HSD, *** p < 0.001, ** p < 0.01, * p < 0.05).

**Fig. S4.**
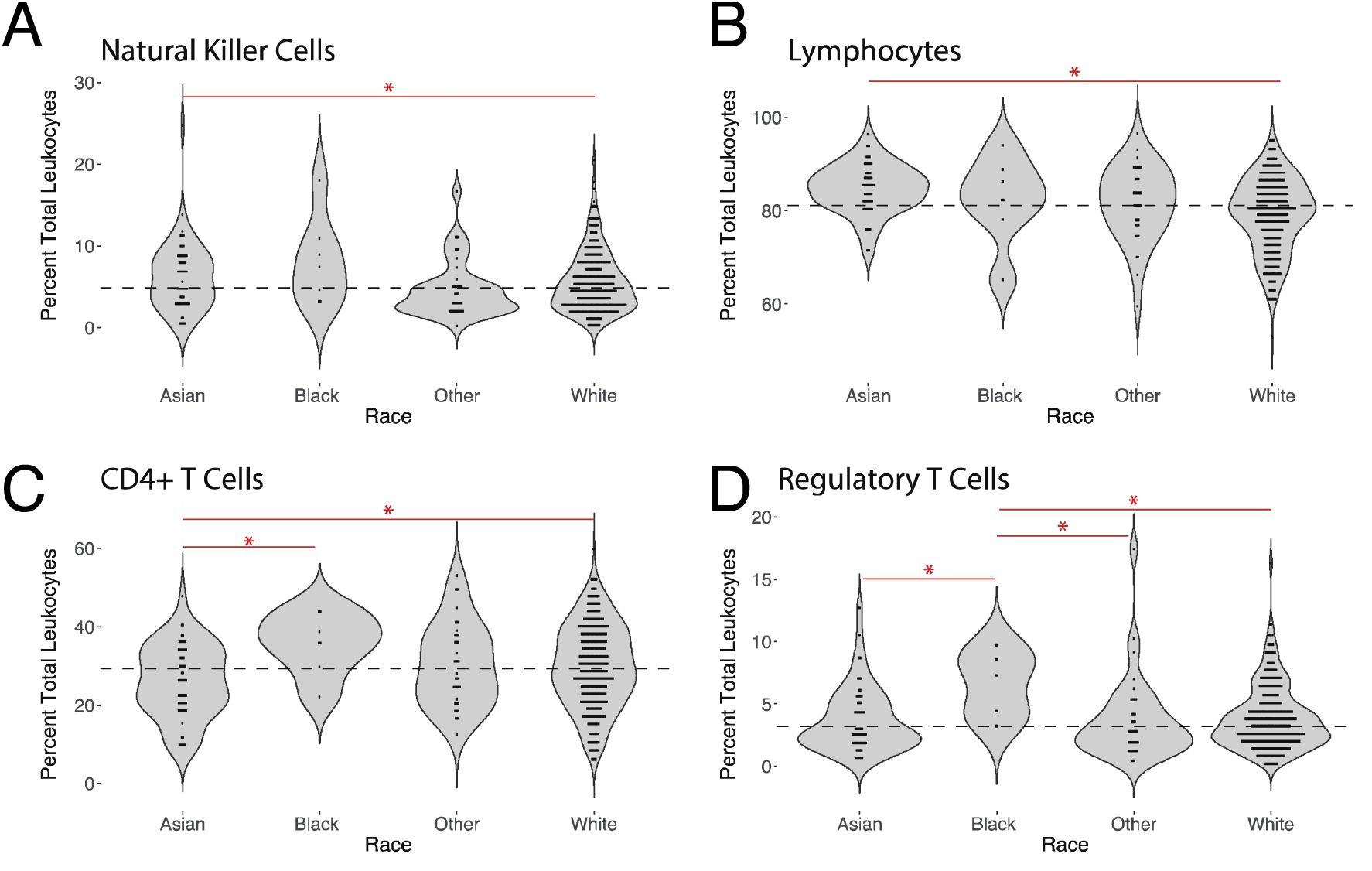
Cell subset measurements differ by race. **(A)** Natural Killer Cells are significantly elevated in Asian, as compared to white subjects, accounting for age and sex (ANCOVA with Tukey HSD, n = 578, * p < 0.05). **(B)** Lymphocytes are significantly elevated in Asian as compared to white subjects, accounting for age and sex (ANCOVA with Tukey HSD, n = 578, * p < 0.05). **(C)** CD4^+^ T cells are significantly decreased in Asian as compared to African American or white subjects, accounting for age and sex (ANCOVA with Tukey HSD, n = 578, * p < 0.05). **(D)** Regulatory T cells are significantly decreased in African American subjects as compared to all other ethnic groups, accounting for age and sex (ANCOVA with Tukey HSD, n = 578, * p < 0.05).

**Fig. S5.**
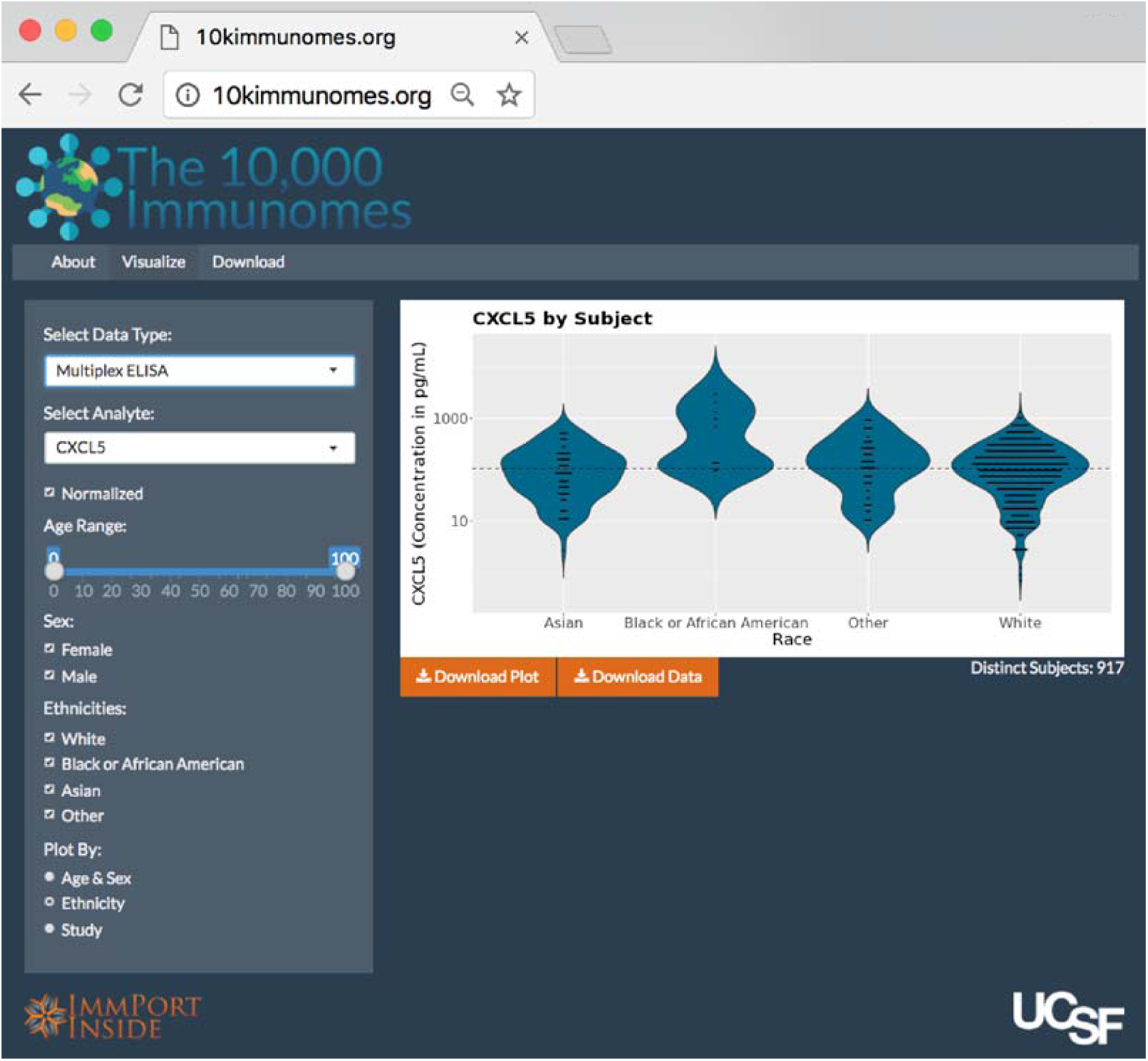
Web Interface. Screen capture of the interactive visualization portion of the web interface, depicting a plot of CXCL5 concentration by race. From this page, users can plot data from the resource by age, sex, race, and study. They can subset which data are plotted based on the demographic variables of their choosing, and they can download the plots as PDF files as well as the underlying data in a standardized format that includes subjects’ demographic information and the study from which the data were obtained.

**Table S1.**
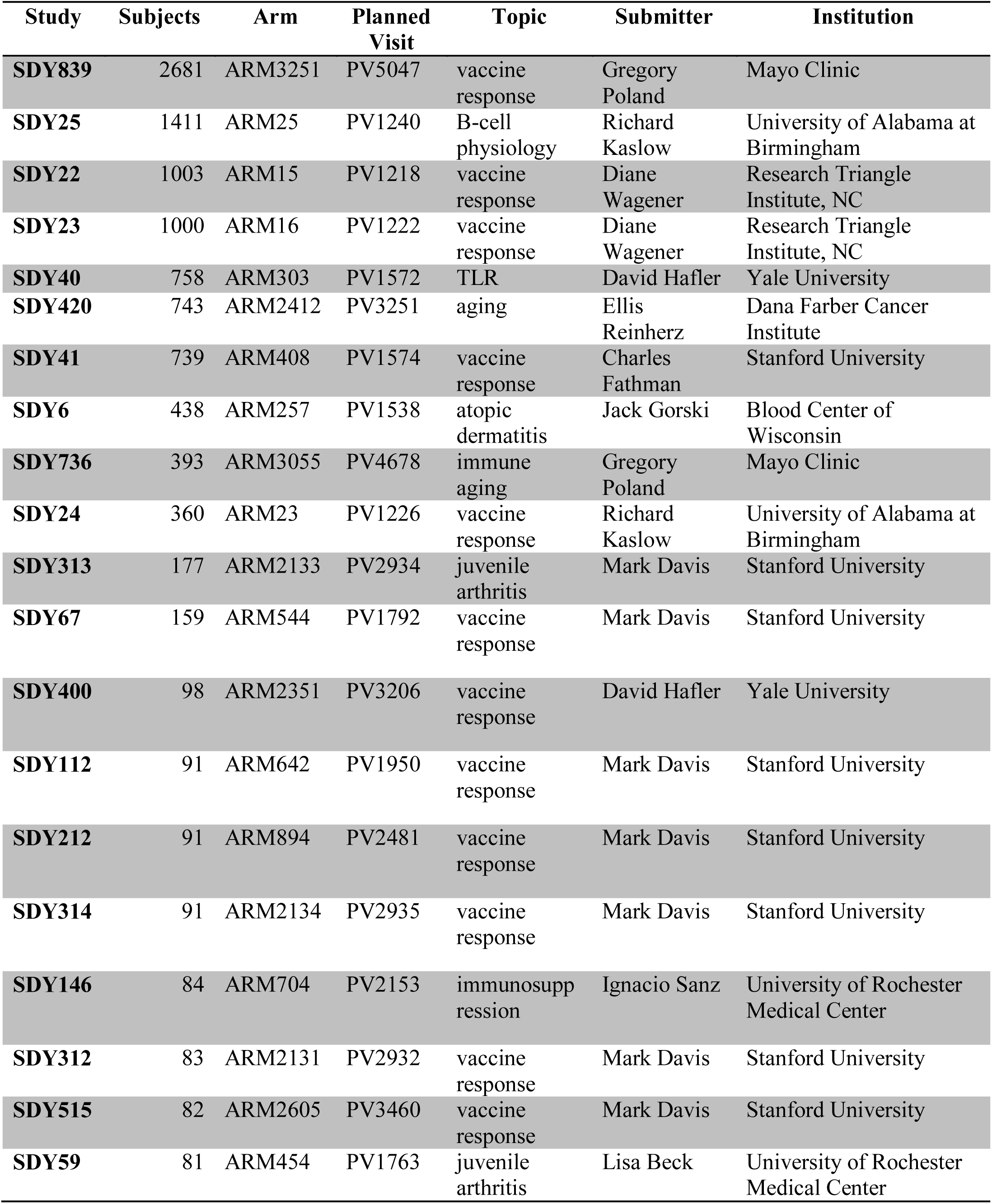
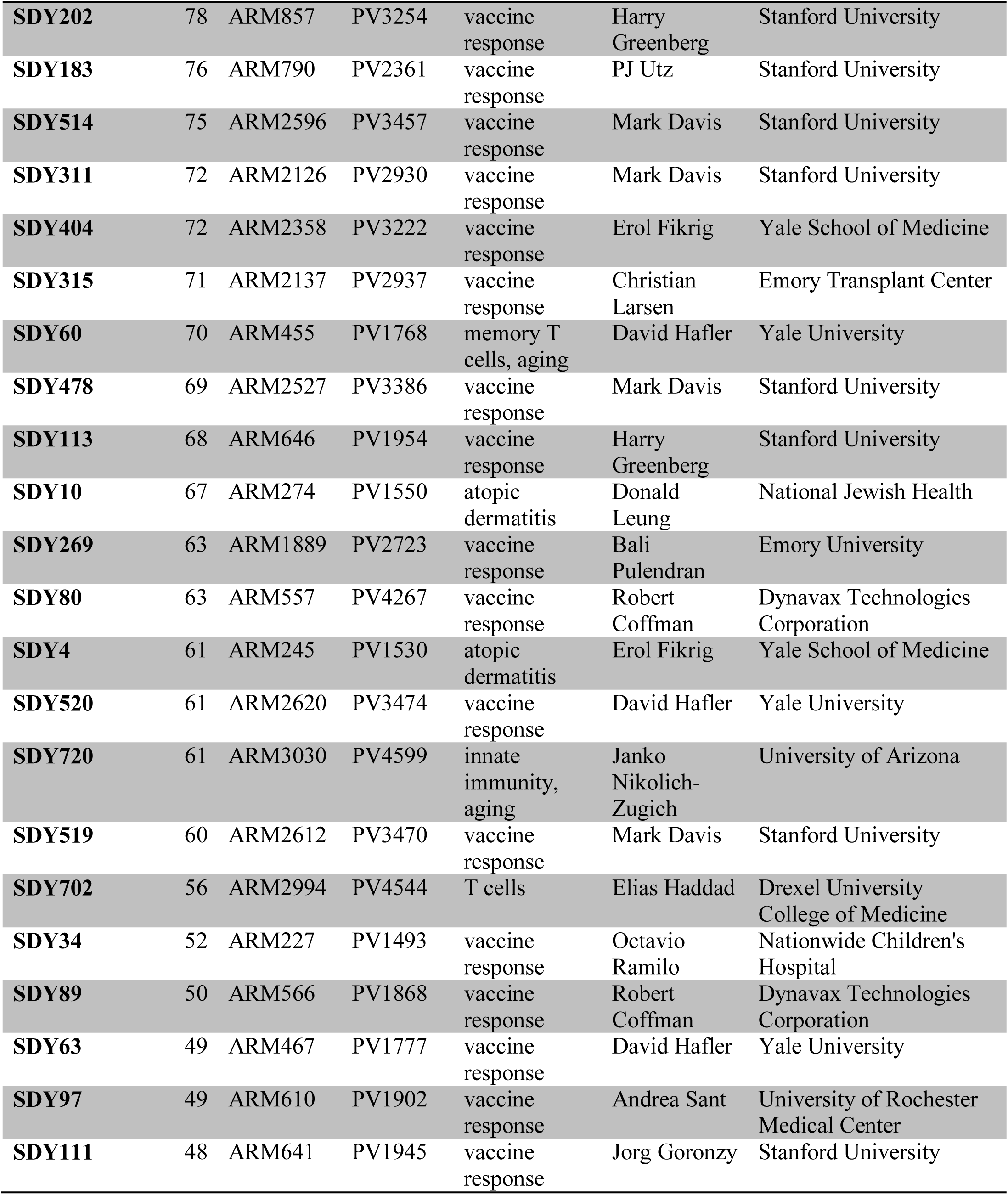
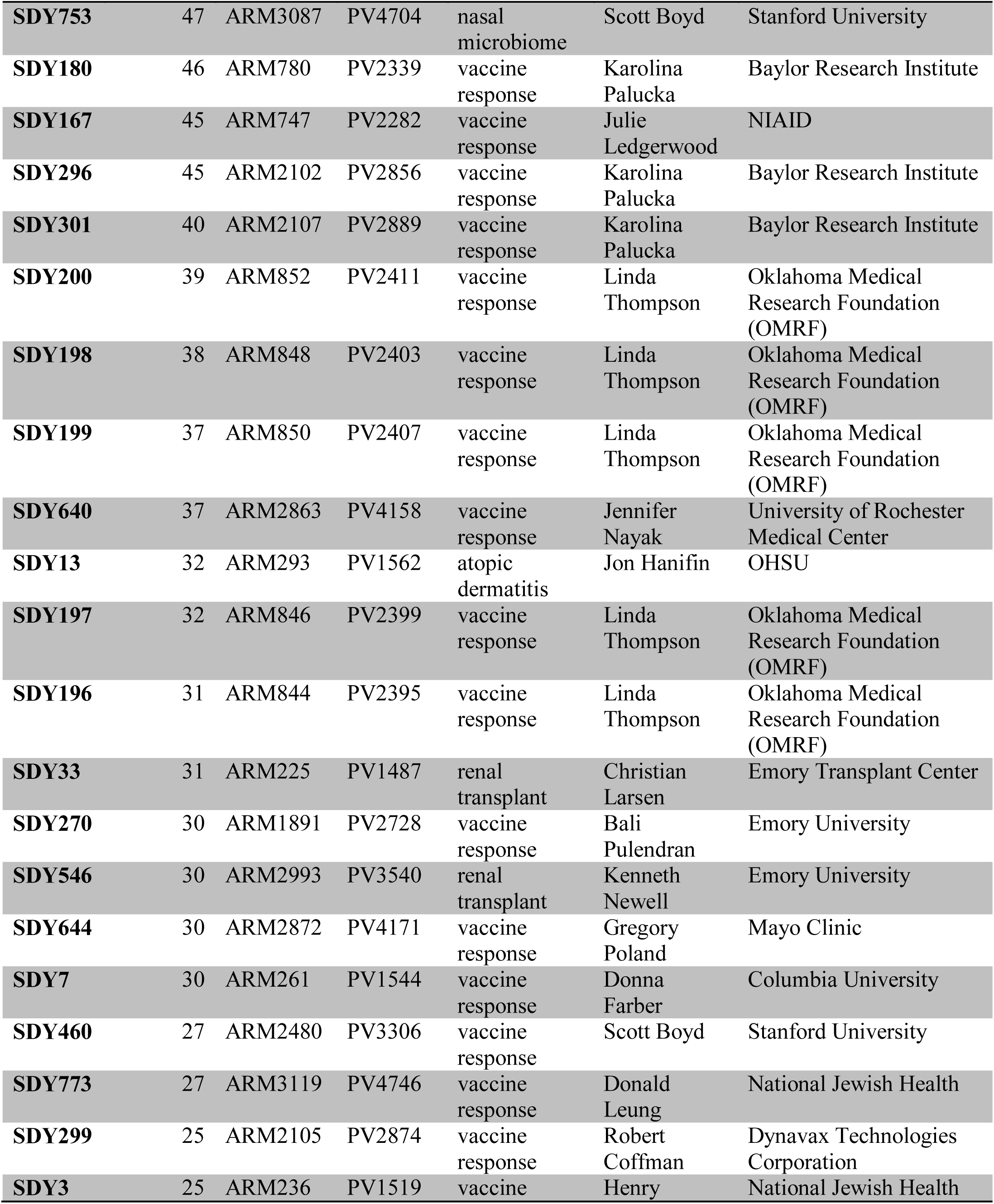
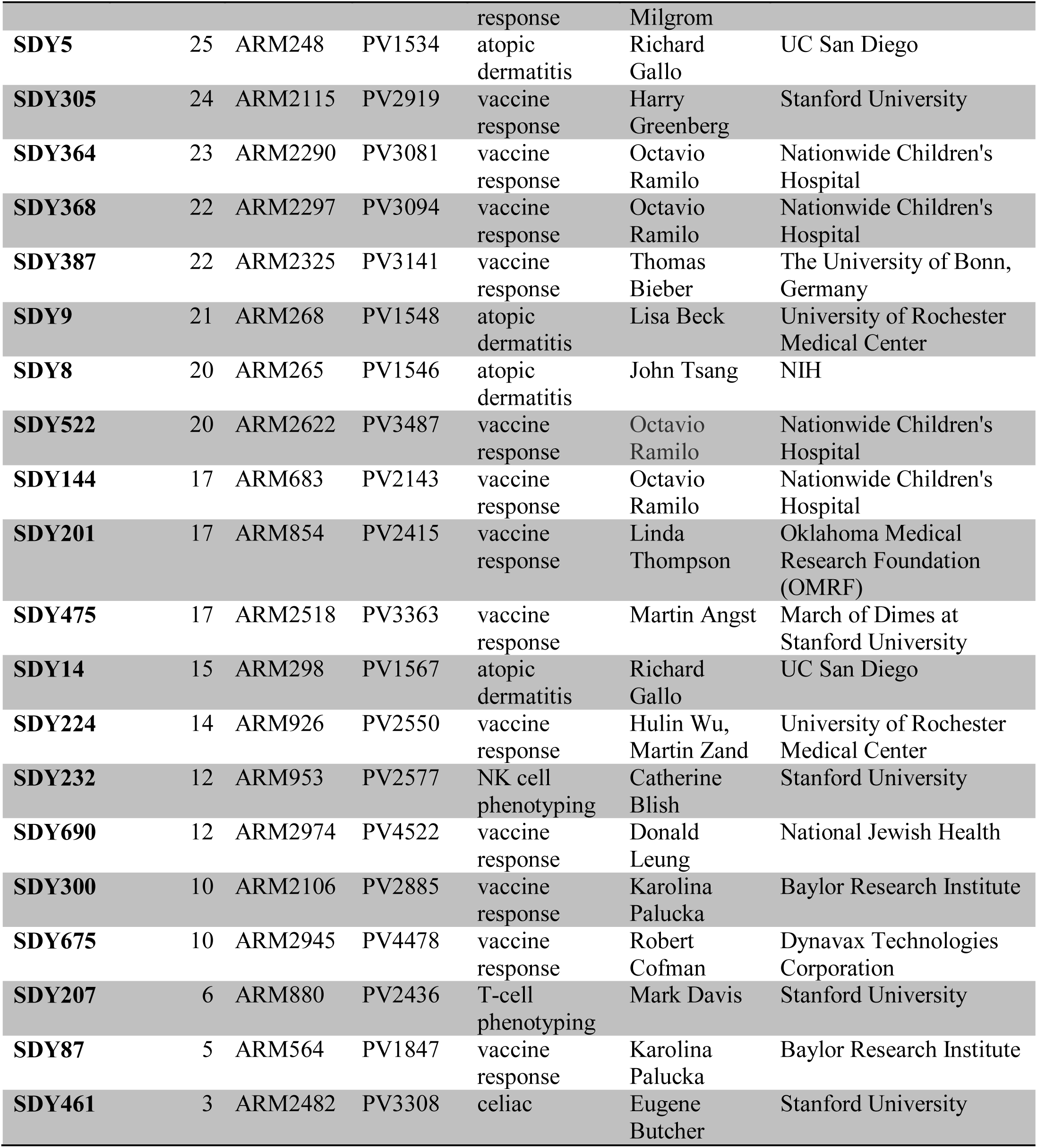
Data included in the 10,000 Immunomes Project. Enumerates the studies, experimental arms, time points (planned visits) determined by manual review to contain samples from healthy control subjects and included in the 10KIP. Each study is associated with its data contributor and their respective institution.

